# Reproducibility problem with a proposed standard method to measure disinfection efficacy

**DOI:** 10.1101/2022.03.15.484496

**Authors:** John Hilgren, Mrudula Srikanth, Milady Brutofsky, Keith Mainquist, Paul Prew, William King, Rhonda Jones, Lisa Hellickson, Josh Luedtke, Kristie Restrepo, Florian Brill, Patrick Quinn, the Efficacy Working Group

## Abstract

A collaborative study was carried out in four laboratories to determine the reproducibility of a proposed ASTM International standard method for quantitatively evaluating the efficacy of disinfectants on hard, non-porous surfaces against bacteria. The method, known as the Quantitative Method, has also been suggested as a future regulatory standard for the United States and internationally. The multi-lab study was carried out using *Pseudomonas aeruginosa* and an alkyl dimethyl benzyl ammonium chloride antimicrobial product diluted in hard water. Results of the study showed acceptable repeatability in log_10_ reductions within each laboratory, but unacceptable reproducibility across laboratories despite careful analyst training and standardization of test conditions. A follow-up study ruled out analyst-to-analyst differences as the cause of the poor reproducibility. As it currently exists, the Quantitative Method is not sufficiently reproducible. Ruggedness testing to assess the sensitivity of the method to small changes in operational factors is recommended.

## Introduction

Reproducibility is critically important to the scientific method, and scientists are increasingly concerned about a “replication crisis” and its impacts on public confidence in science [1]. The disinfectant industry and regulators in the United States (US) have been on a multi-decade journey to address this challenge. This US industry’s journey began in 1990 when concerns about the efficacy of household disinfectants led the US House Committee on Government Operations to request that the General Accounting Office (GAO) review the Environmental Protection Agency’s regulation of these products [2]. The GAO reported that the US *gold standard* to measure disinfection efficacy – the AOAC Use-Dilution Method (UDM) - may not be reproducible. Industry and regulators responded by developing a new method - the AOAC hard surface carrier test (HSCT) [3]. The HSCT was at one time accepted by the Environmental Protection Agency (EPA) to support the registration of liquid disinfectants; however, interlaboratory studies to validate the method with soil and hard water were never completed so it was not embraced by users and is no longer included in EPA’s 810.2000 guidance for efficacy testing to support product registration [4].

By 2002, increasing interest in global harmonization of disinfectant test methods led to an international workshop on disinfectant test methods [5]. The workshop included a review of the pros and cons of two existing standards: Committee of European Norms EN 13697 [6] and ASTM International Standard E2197 [7]. One outcome from the workshop was that the Organization of Economic and Cooperative Development (OECD) funded a scientific review of disinfectant test methodologies. The review gave rise to a specification for technical elements to include in a future international method [8]. Subsequently, the Quantitative Carrier Tests (QCTs) ASTM E2111, ASTM E2197, and EN 13697 were evaluated as platforms upon which to build a future OECD method. In the end, ASTM E2197 was selected as the optimal candidate [9, 10, 11]. Research to develop an OECD version of E2197 was carried-out from 2007 to 2009; thirty-five laboratories from eight OECD member countries participated [12]. In 2009, the U.S. EPA issued their own guidance for testing disinfectant efficacy against *Clostridioides difficile* spores using a QCT method based on ASTM E2197 [13]. The EPA drafted a QCT method in 2010 for potential testing of disinfectants against vegetative bacteria [14]. The method, MB-25 version 00, was also based on ASTM E2197. In 2011, results from the OECD member country research were presented to the OECD Working Group of National Coordinators (WNT) along with a request to leverage the work to create an OECD Test Guideline. The WNT rejected the proposal because of multiple technical concerns. In 2011, an *ad hoc* group of experts began collaborating to address WNT’s concerns. The group’s work succeeded in gaining WNT’s support to publish four QCT methods in an OECD guidance document [15]. The four QCTs measure bactericidal, mycobactericidal, fungicidal and virucidal efficacy. Meanwhile, also in 2011, the Efficacy Working Group (EWG), a consortium of individuals representing disinfectant ingredient suppliers and producers, met with EPA and OECD where, among other things, they expressed concerns about the reproducibility and bias of the draft OECD method. In 2012, the EPA issued MB-25 version 01 which changed the broth medium used to cultivate *Pseudomonas aeruginosa* from Tryptic Soy Broth (TSB) to Synthetic Broth (SB); it also added new instructions on carrier manufacturing. The OECD guidance changed from TSB to SB in 2013. In 2017, the EPA issued guidance for using a QCT method for testing efficacy against *Candida auris* [16].

In 2018, the EWG raised three concerns to EPA about further implementation of QCT methods to support disinfectant product registrations: variations in carrier quality, poor method reproducibility, and result bias relative to the UDM *gold standard.* In response to these concerns, the EWG and EPA began collaborative research. This research led to implementation of a new protocol for preparing and sterilizing the soil load (a mixture of bovine serum albumin, mucin, and yeast extract) and new quality specifications for carriers. Instead of updating MB-25 with these changes, EPA introduced them in a new method named, *“Test Method for Quantitatively Evaluating the Efficacy of Antimicrobial Products on Hard, Non-Porous Surfaces Against Bacteria”,* also referred to as the Quantitative Method (QM). In 2020, when comparing the results from the QM to the UDM, the EWG observed greater than 2-log_10_ differences in disinfection results across laboratories when testing with *P. aeruginosa* and n-alkyl dimethyl benzyl ammonium chloride (ADBAC; Barquat® MB-50, Arxada, Basel, Switzerland) diluted in hard water. The log reduction differences caused EWG to halt research on QM versus UDM bias and shift their focus to QM reproducibility. It should be noted that efficacy against *P*. *aeruginosa* is of practical importance for two reasons: *P. aeruginosa* is used by the EPA to differentiate Hospital-level disinfection from Broad-Spectrum disinfection, and ADBAC disinfectants are used widely around the world for disinfection of hard surfaces including those in hospitals and healthcare facilities [17]. After finding no material or method issues that could explain the lab-to-lab differences, the EWG laboratories embarked on ruggedness studies to assess the sensitivity of the QM to small changes in a variety of different operational factors. This ruggedness testing revealed that efficacy of ADBAC against *P. aeruginosa* is highly sensitive to the concentration of 3-part soil, and more specifically, the concentration of the mucin component.

In August 2021, the U.S. EPA initiated a working group within ASTM International to evaluate the QM for adoption as an ASTM Standard. Within ASTM, the EPA shared data generated in their laboratory on the responsiveness of the QM when testing an ADBAC product (BTC® 835, Stepan Company, Northbrook, IL) in hard water against *P. aeruginosa.* Based on the results of their study, the EPA suggested the ADBAC concentrations and exposure times used in their study be considered for use as a test system or proficiency control in the QM. Three EWG member laboratories, and one independent laboratory in Germany, using the QM provided by the EPA, then replicated the EPA study resulting in the data included in this manuscript. The data in this manuscript confirms a reproducibility problem with the QM. Before proceeding with ASTM interlaboratory studies to measure method performance (which assumes expert concurrence of method readiness and leads to development of regulatory standards), we suggest additional method ruggedness research. Investing in this research should help avoid two problems of the past: implementation of a method not used by the community it is intended to serve, and implementation of a method that raises public concern about the reliability of disinfectants. A reproducible QM should bolster adoption as an ASTM standard and as a harmonized global standard for disinfection.

## Materials and methods

### Bacterial test culture preparation

The bacterial strain used throughout this study was *P. aeruginosa* ATCC® 15442 (American Type Culture Collection, Manassas, VA). The bacterial test culture was prepared following the QM [19]. In brief, 100 μL of a thawed single use frozen stock culture was used to inoculate 10 mL of Synthetic broth with 10% dextrose (w/v) added just prior to use. After incubating at 36°C for 24 h, the pellicle was removed from the culture. The broth culture just below the removed pellicle was carefully collected avoiding any sediment at the bottom of tube and transferred to a 15 mL conical centrifuge tube. This broth culture was then centrifuged at 5,000 gN for 20 minutes. The supernatant was removed without disrupting the pellet, then the pellet was resuspended in 6.5 to 10.0 mL phosphate buffered saline (PBS) resulting in a *P. aeruginosa* test suspension that was ready for soil addition. The amount of PBS added was based on prior testing experience to achieve a mean density of bacteria on dried control carriers of 5.0 to 6.0 log_10_ colony forming units (CFU) per carrier.

### Soil load and final test suspension preparation

A three-part soil was prepared according to the QM [19]. In brief, three stock components were prepared separately; (i) 0.5 g of bovine serum albumin was dissolved in 10 mL of PBS, (ii) 0.5 g of yeast extract was completely dissolved in 10 mL of PBS, and (iii) 0.04 g of bovine mucin was dissolved in 10 mL of PBS. After dissolution in PBS, each stock solution was filter-sterilized by passing through a 0.2 μm pore size (polyethersulfone) membrane filter. The sterilized stock solutions were then aliquoted into single use quantities and stored at −20°C until use. On the day of testing, 160 μL of a three-part soil mixture (comprising 25 μL BSA, 35 μL yeast extract and 100 μL bovine mucin) was added to 340 μL of the *P. aeruginosa* test suspension. The final test suspension with soil was mixed and used to inoculate carriers within thirty minutes.

### Disinfectant working solution preparation

Working solutions containing 500 parts per million (ppm) and 1500 ppm of ADBAC (BTC® 835, 50% n-alkyl dimethyl benzyl ammonium chloride, Stepan Company, Northbrook, IL), were used in the study. The ADBAC test substance was dispensed using a positive displacement pipette. The disinfectant working solutions were prepared in 375 ppm hard water according to the QM [19] and in alignment with EPA 810.2000 guidance for testing disinfectant concentrates. Working solutions were used within three hours of preparation.

### Efficacy testing procedure - repeatability and reproducibility study

Efficacy testing was performed according to the QM [19] in four separate laboratories: A, B, C and D. The analysts in each laboratory were trained, and had prior experience, in running the QM. A brief description of the testing procedure follows.

A summary of the study design is provided in Table 1. Sterile carriers (1 cm diameter discs made of American Iron and Steel Institute Type 304 Stainless Steel with 150 grit unidirectional finish on one side) were inoculated with 10 μL of the final test organism suspension with soil using a positive displacement micropipette. The inoculated carriers, in an open Petri dish, were transferred to a vacuum desiccator and held under vacuum (0.068-0.085 MPa) at room temperature (approximately 22°C) for 45-60 minutes. The carriers were inspected to verify that the inoculated spots were visually dry before being removed from the desiccation unit. The carriers were individually transferred with the inoculated side up to flat-bottom vials to prepare them for disinfectant or PBS control exposure. The exposure was carried out by adding 50 μL of a disinfectant working solution (500 or 1500 ppm ADBAC) or 50 μL of PBS on the carrier to completely cover the dried inoculum spot. At the end of the exposure time, 10 mL of a neutralizing agent, Letheen broth with 0.5% Tween® 80 and 0.07% Lecithin, was added to each vial at timed intervals. The vials were capped, and then vortex mixed to suspend bacteria in the neutralizing solution. The contents of each vial, and 10-fold serial dilutions of each vial in PBS, were filtered using pre-wet membrane filters (0.2 μm pore size, polyethersulfone). The membrane filters were deposited on Tryptic soy agar and incubated at 36°C for 48 h (control carriers) or 72 h (treated carriers) per the QM. Growth on filters was confirmed to be *P. aeruginosa* based on Gram staining and examination for typical colony characteristics. Colonies were enumerated and used for calculation of log_10_ reductions.

**Table 1.**
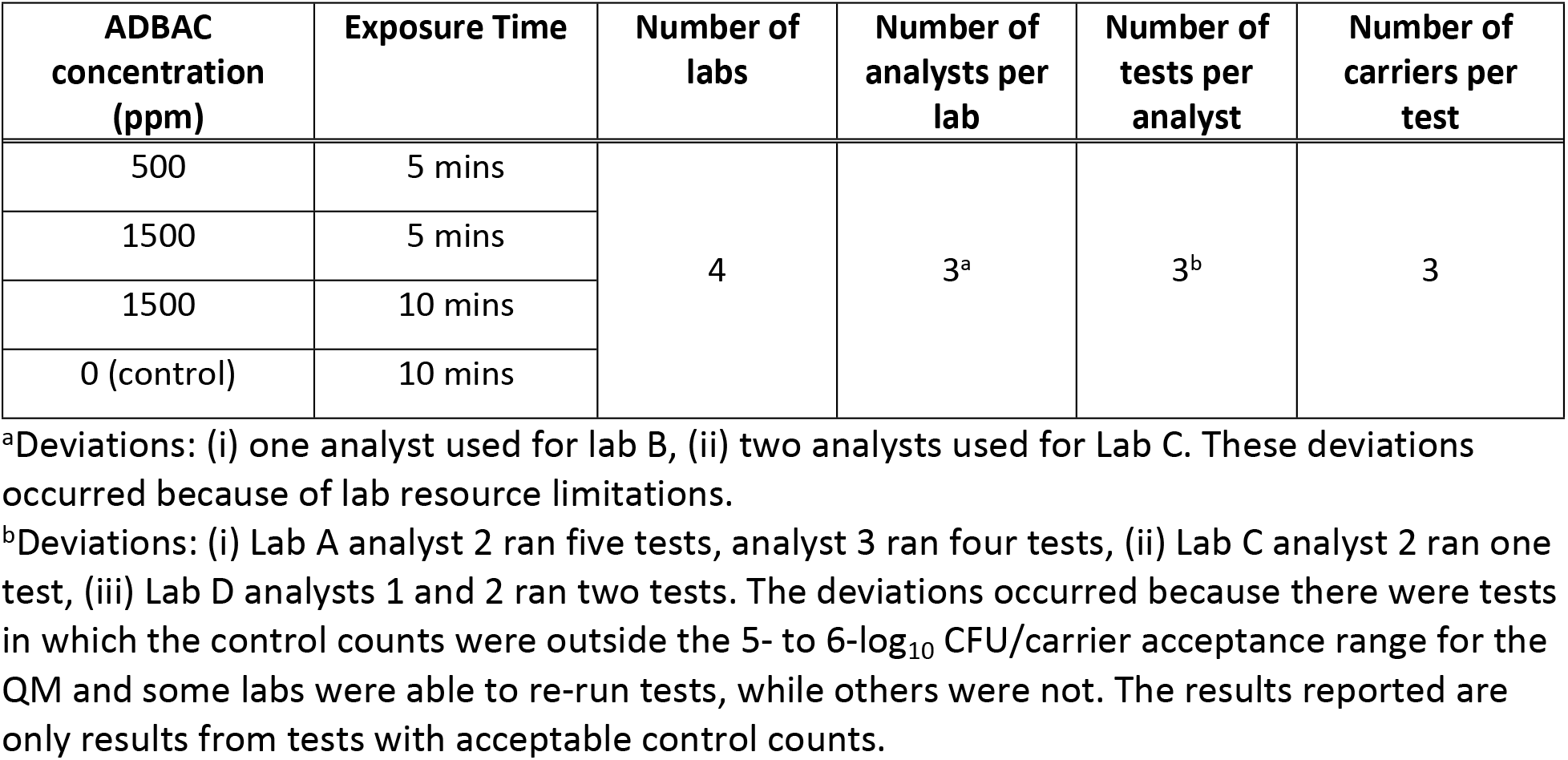
Repeatability and reproducibility study design.

Data were analyzed for repeatability, σ_r_, and reproducibility, σ_R_. Following the convention of Tilt [18] meaning that repeatability is the within-lab precision of log_10_ reductions and includes contributions from analyst and test number variations. Reproducibility is equated with the total precision of log_10_ reductions and includes both within-lab and lab-to-lab variances. The log_10_ reduction is equal to N_C_-N_t_, where N_t_ is the log_10_ of the treatment count and N_C_ is the log_10_ of the corresponding control count. Test number is nested within analyst and analyst within lab, both as random effects. The data were analyzed using JMP® statistics software (Cary, NC). Because the data was unbalanced, the Restricted Maximum Likelihood procedure was employed. Each treatment condition was analyzed separately.

### Efficacy testing procedure - analyst effects study

Analysts from Labs A, C and D from the repeatability and reproducibility study gathered at the EPA Environmental Science Center laboratory (Fort Meade, MD) on the same day to test analyst-specific effects on log_10_ reductions.

Efficacy testing was performed according to the QM [19]. All analysts used materials provided by the EPA laboratory. Labs A and C had 2 persons each working as a team and Lab D had 1 person only. The ADBAC treatments and testing procedures used in this study were the same as those used in the repeatability and reproducibility study. Each analyst team prepared their own set of control carriers and tested three carriers per treatment. Data analysis of repeatability was carried out in the same manner as the earlier study.

In this study, there were no replicates within each analyst team. For each treatment condition the statistic of interest is the standard deviation of log_10_ reduction across the analyst teams, σ_AT_. This metric can be viewed as either a repeatability measure, since it is based on the within-lab (or within-facility or within-location) variance across analyst teams performing the tests in that one facility, or it can be viewed as an estimate of reproducibility since it represents the company-to-company variance made up of the same company analyst teams as in the first study.

## Results

### Repeatability and reproducibility study

Figure 1 displays the control counts (N_c_) across all the tests and Table 2 provides a summary of the statistical analysis results, which show excellent repeatability (σ_r_=0.19) and reproducibility (σ_R_=0.25). As noted previously, there were multiple tests in which the control counts were outside of the acceptable range, and results from those invalid tests are not included in Table 2 or elsewhere in this paper. This highlights a need for a closer look at the final test suspension preparation procedures to reduce the number of invalid tests.

**Fig 1.**
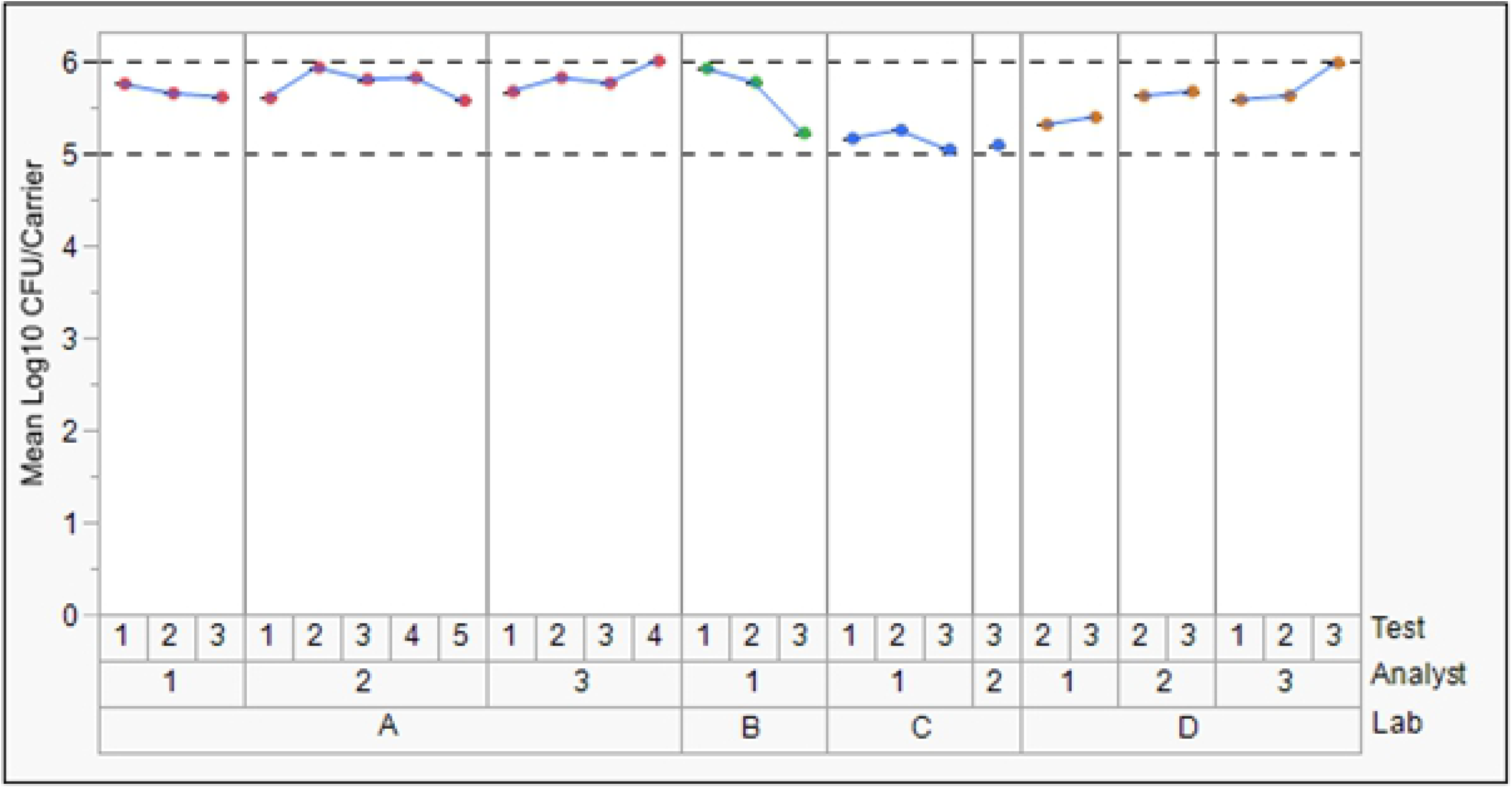
*P. aeruginosa* untreated controls. Markers are the average of three carriers, dotted lines are the low and high acceptance bounds.

**Table 2.** Repeatability and reproducibility of *P. aeruginosa* control and treatments.

| ADBAC concentration (ppm) | Exposure time (min) | Repeatability<sup>a</sup><br>Standard Deviation<br>($\sigma_r$) | Reproducibility<sup>b</sup><br>Standard Deviation<br>($\sigma_R$) |
| --- | --- | --- | --- |
| 0 (control) | 10 | 0.19 | 0.25 |
| 500 | 5 | 0.25 | 1.27 |
| 1500 | 5 | 0.60 | 1.62 |
| 1500 | 10 | 0.70 | 0.93 |

<sup>a</sup>Repeatability is the within-lab precision of $\log_{10}$ reductions and includes contributions from analyst and test number variations
<sup>b</sup>Reproducibility is equated with the total precision of $\log_{10}$ reductions and includes both within-lab and lab-to-lab variances

Table 3 lists the log_10_ reductions of all the tests for the three separate treatment conditions by laboratory. Figures 2, 3 and 4 provide additional details on the log_10_ reductions by displaying the results by individual analyst and test within each laboratory. The three treatment conditions captured an acceptably wide efficacy range. A summary of repeatability and reproducibility of the log_10_ reductions is shown in Table 2. The repeatability measures are 0.25, 0.60, and 0.71 for the three treatment conditions, and the reproducibility measures are 1.27, 1.62, and 0.93 for the same respective treatment conditions. For log_10_ reductions in the midrange, repeatability standard deviations trend higher as can be seen in Figures 3 and 4. Of greater concern are the reproducibility statistics for the low and medium efficacy conditions (500 ppm at 5 minutes and 1500 ppm at 5 minutes) for which the first of these has a reproducibility of σ_R_=1.27, which is close to the mean log_10_ reduction, and for the second of these, a σ_R_=1.62 which exceeds proposed acceptance thresholds included in the discussion.

**Table 3.** Reductions of *P. aeruginosa* by laboratory.

| ABDAC concentration (ppm) | Exposure time (min) | Mean Log<sub>10</sub> Reduction |  |  |  |
| --- | --- | --- | --- | --- | --- |
|  |  | Lab A | Lab B | Lab C | Lab D |
| 500 | 5 | 0.35 | 0.98 | 3.09 | 0.58 |
| 1500 | 5 | 1.81 | 1.85 | 4.96 | 1.74 |
| 1500 | 10 | 3.57 | 3.81 | 5.14 | 3.98 |

**Fig 2.**
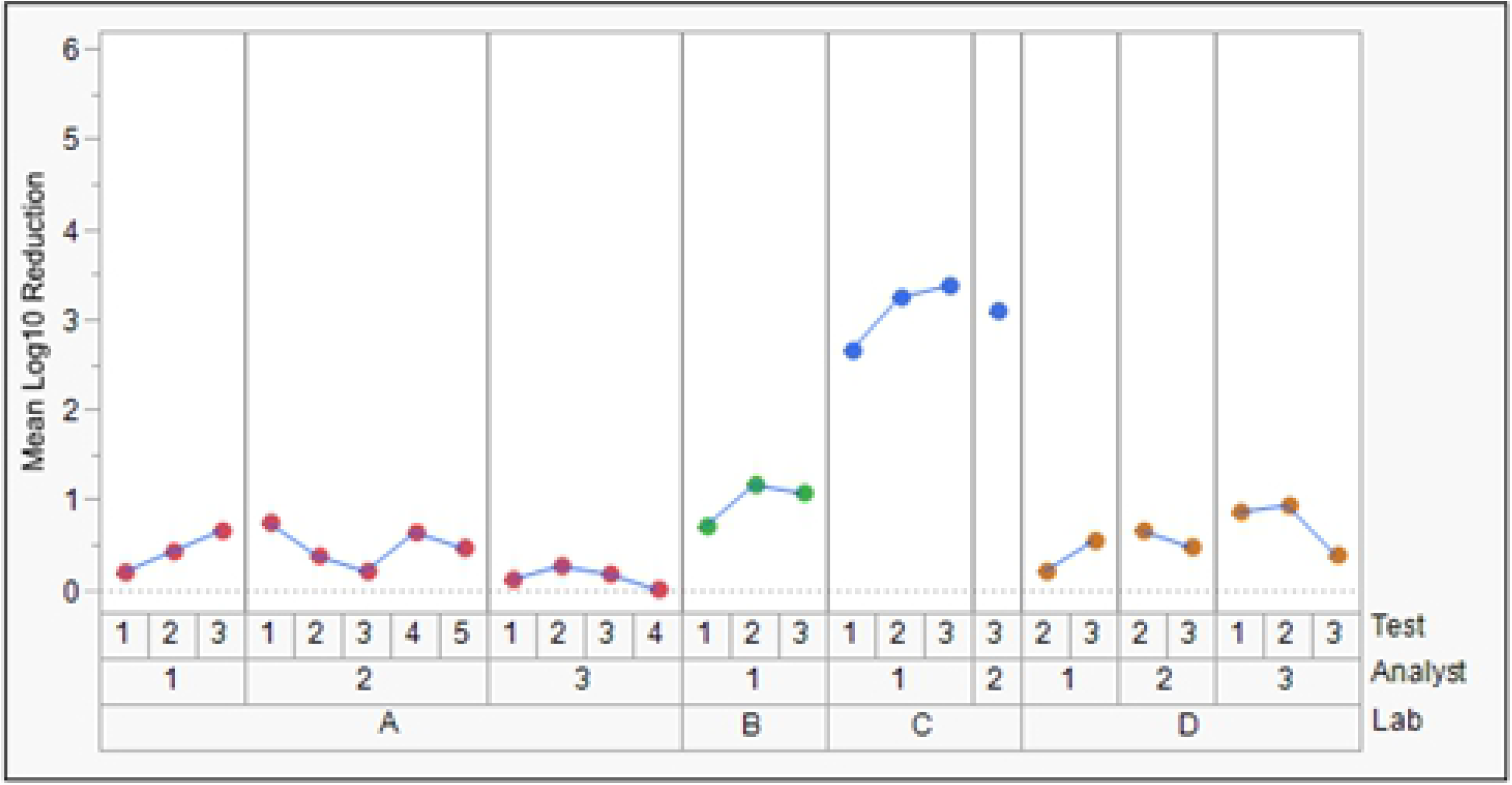
Reductions of *P. aeruginosa* exposed to 500 ppm ADBAC for 5 minutes. Markers are the average of 3 carriers.

**Fig 3.**
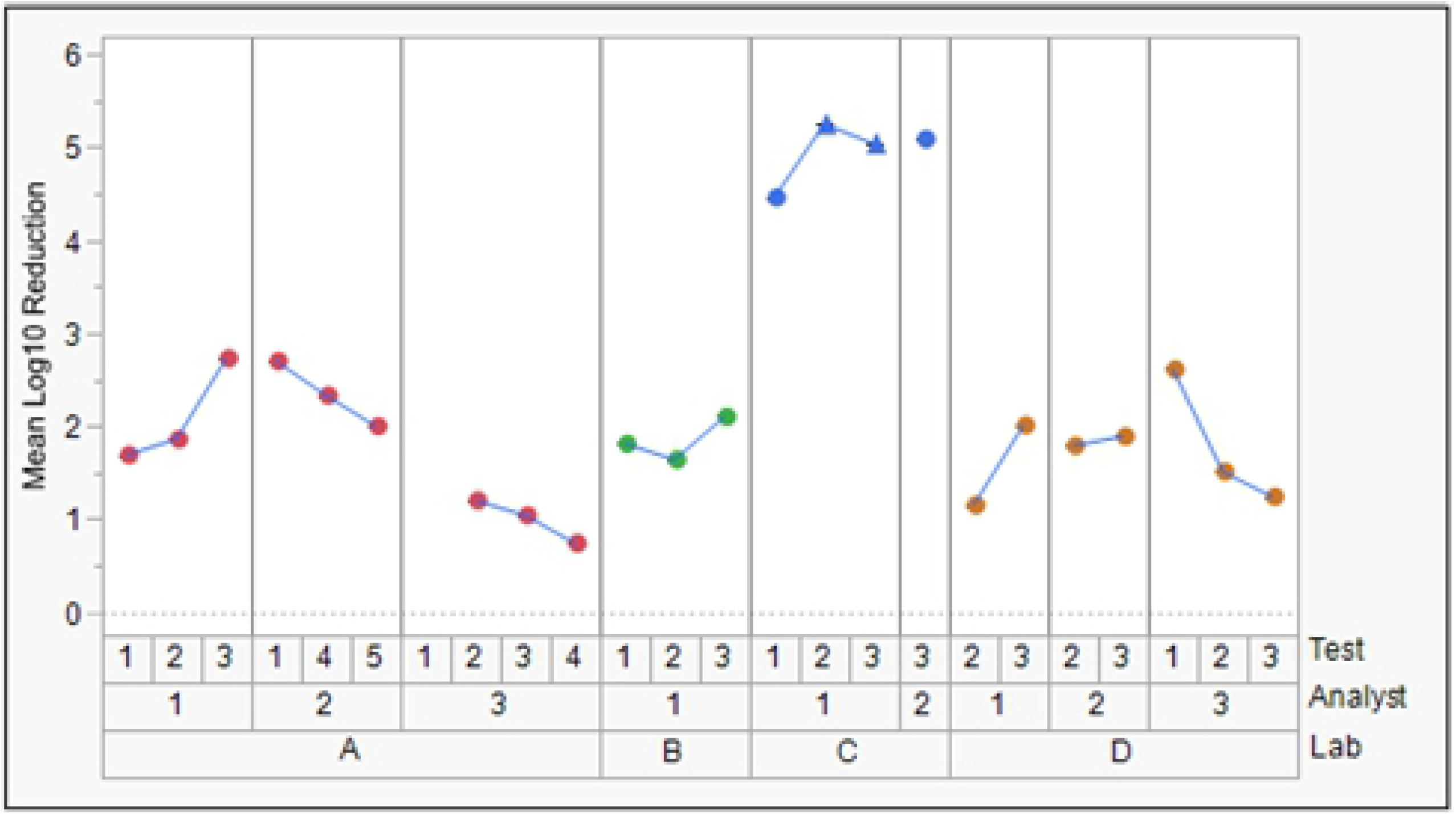
Reductions of *P. aeruginosa* exposed to 1500 ppm ADBAC for 5 minutes. Markers are the average of 3 carriers, and triangles represent complete kill.

**Fig 4.**
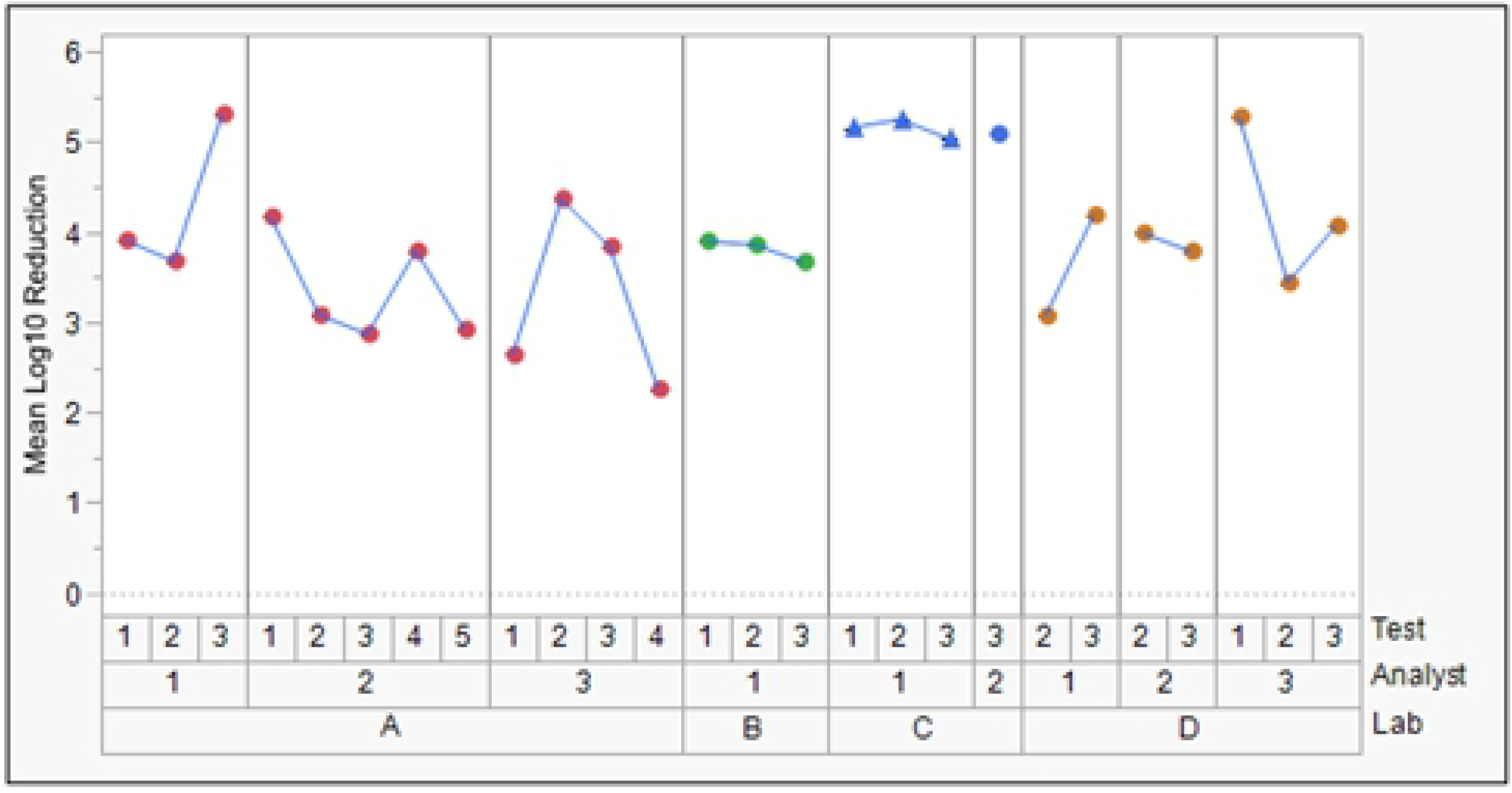
Reductions of *P. aeruginosa* exposed to 1500 ppm ADBAC for 10 minutes. Markers are the average of 3 carriers, and triangles represent complete kill.

### Analyst effects study

Figure 5 is a plot of the control counts (N_C_’s) across the three analyst teams – all lie within the acceptable range. Figure 6 is a plot of the log_10_ reductions for the three analyst teams for all the treatment conditions. The standard deviations across analyst teams (σ_AT_’s) are provided in Table 4. For all three treatment conditions, σ_AT_ is impressively small (0.13, 0.29, 0.08 for the low, medium and high efficacy treatment conditions). These are in striking contrast to the σ_R_’s of the first study. The 1.78 and 4.24 log_10_ reduction for the low and medium efficacy conditions do not corroborate the results from any of the labs in the repeatability and reproducibility study, further highlighting the concern of poor reproducibility.

**Fig 5.**
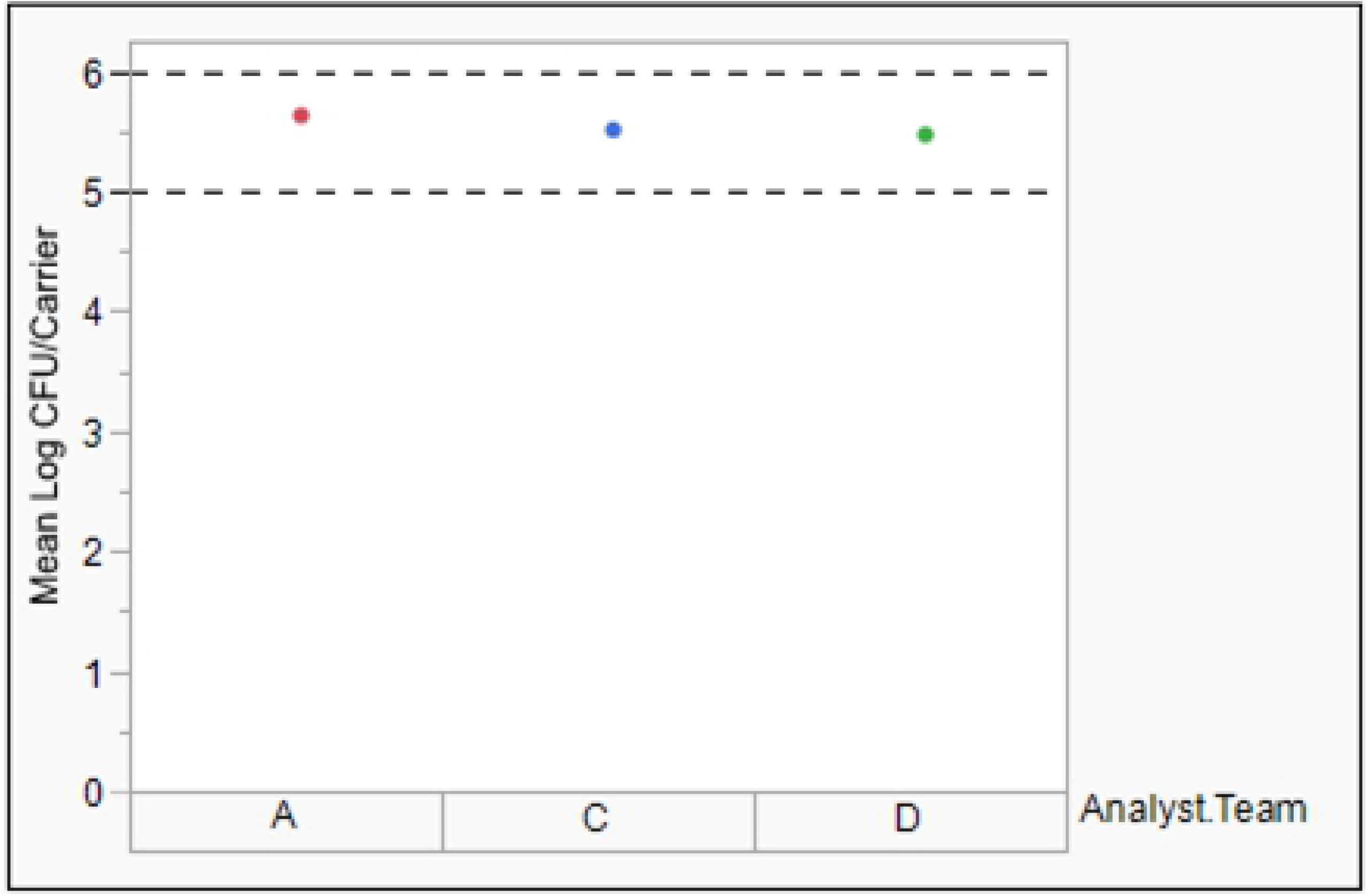
*P. aeruginosa* untreated controls – analyst effects study. Markers are the average of three carriers, and the dotted lines are the low and high acceptance bounds.

**Table 4.** Variance of *P. aeruginosa* results.

| ADBAC concentration (ppm) | Exposure time (min) | Mean Log<sub>10</sub> reduction across Analyst Teams | Standard deviation across Analyst Teams ($\sigma_{AT}$) |
| --- | --- | --- | --- |
| 500 | 5 | 1.78 | 0.13 |
| 1500 | 5 | 4.24 | 0.29 |
| 1500 | 10 | 5.54 | 0.08 |

**Fig 6.**
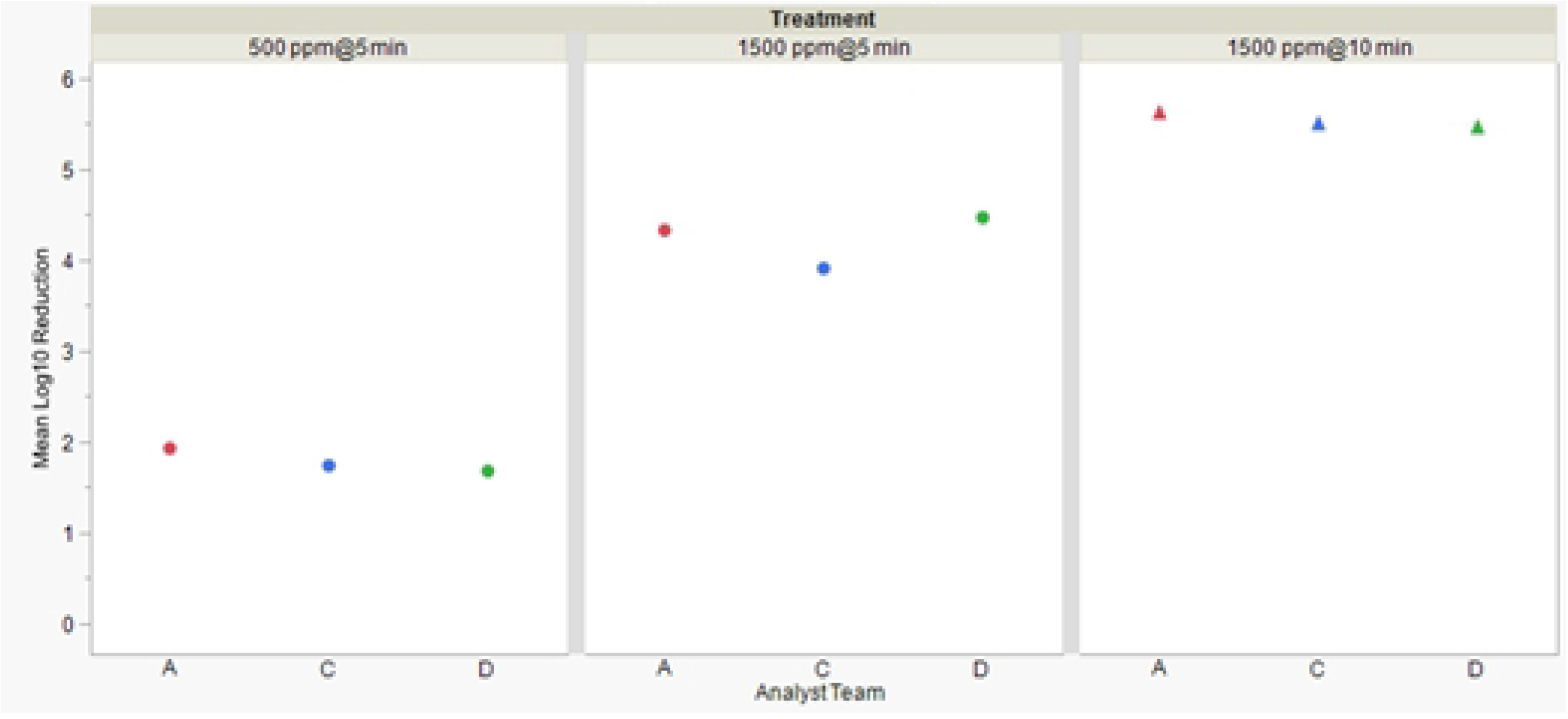
Reductions of *P. aeruginosa* exposed to ADBAC treatments – analyst effects study. Markers are the average of 3 carriers, and triangles represent complete kill.

## Discussion

It is important to understand the repeatability and reproducibility of a test method before deciding on its applicability as a regulatory standard. For antimicrobial efficacy methods, assessments of repeatability and reproducibility are best carried out using treatments delivering a medium level of effectiveness. For example, it is best to use treatments delivering 2- to 4-log_10_ reductions when a method (such as the QM) has an upper limit close to a 5-log_10_ reduction. This is because the results when testing ineffective treatments and highly effective treatments are always more reproducible than those from moderately effective treatments. Currently, little data exists on the ruggedness, repeatability, and reproducibility of the QM at any level of antimicrobial effectiveness despite interest in adopting it as an international standard. Data on the reliability of the QM under medium effectiveness conditions was needed to help technical experts from across government, academia and industry determine appropriate next steps in the development of the QM.

In this study, we measured the efficacy of a quaternary ammonium chloride-based disinfectant against *P. aeruginosa* in four different laboratories using the QM. The selection of a quaternary ammonium chloride-based disinfectant was important because it represented a product group widely used in institutional and healthcare settings. Three different combinations of disinfectant concentration and exposure time were chosen because preliminary data suggested they would result in low, medium, and high numbers of treatment survivors, minimizing no kill or complete kill outcomes which are less useful for a repeatability and reproducibility study.

The data in this paper highlights an obvious reproducibility problem with the QM. Specifically, log_10_ reductions using the medium efficacy treatment (1500 ppm active ingredient with a 5-minute exposure) were approximately 3-log_10_ units higher in one of the laboratories than the others despite careful standardization of materials, methods, and training in all laboratories prior to the study. Higher log_10_ reductions were also observed in the same laboratory with the other treatments; however, the magnitudes of the differences were less, as expected. Interestingly, the QM results from the laboratory with the higher log_10_ reductions correlate closely with historical UDM results whereas those from the other labs do not (unpublished EWG data). The degree of correlation between QM and UDM results is an important consideration because it has a direct impact on how a disinfect product is applied. For example, if the QM is harder to pass than the UDM, then products developed and registered with the QM would need to be used at a higher active ingredient concentration or for a longer exposure time. A risk-benefit analysis considering the impacts of disinfectant application changes on public health and the environment would help set acceptance limits for test method correlation.

The reproducibility standard deviation for the medium efficacy treatment in this collaborative study was 1.62 log_10_. A knowledge sharing article published by Montana State University (MSU) recommends using an acceptability criterion of <1.5 log_10_ for deciding whether the reproducibility standard deviation of a disinfectant test method is acceptable [20]. A limitation of MSU’s criterion is that it is based on historical precedent rather than microbial risk assessment or another evidence-based evaluation. In response to the need for an improved process for making decisions about test method reproducibility, Parker published a process for assessing antimicrobial test method reproducibility based on stakeholder’s specifications [21]. Parker’s process requires a stakeholder to specify the percentage (γ) of the tests that must produce log reductions that differ from an ideal true log_10_ reduction value (μ) by no more than a maximum error (δ). The stakeholder specifications, coupled with data from a multi-laboratory study of an antimicrobial test method, can then be used to determine how small the reproducibility standard deviation must be to conclude that the method is acceptably reproducible. We suggest that the ASTM International QM working group use Parker’s process, along with the data in this publication, to determine if additional research is warranted before embarking on larger interlaboratory studies. We propose using the following values for μ, δ, and y:

1. Set the ideal true log_10_ reduction (μ) to 4.0. The ideal true log reduction serves as the *de facto* regulatory performance standard. Quantitative microbial risk assessment (QMRA) models by Ryan [22] predicted that a 2-log_10_ bacterial reduction, and a 3-log_10_ virus reduction, on fomites could reduce infection risk from a single fomite contact to less than one-in-a-million. Later studies by Wilson [23] predicted that a 4-log_10_ reduction of rotavirus and rhinovirus would be needed to achieve a one-in-a-million risk target when there is six hours of fomite contact activity.
2. Set the maximum error (δ) to 1-log_10_. If the ideal true log_10_ reduction is 4.0, then a δ of 1-log_10_ translates to a range of 3- to 5-log_10_ reduction which still maintains a reasonable level of infection risk reduction based on the QMRA studies referenced above.
3. Use 95% for the percentage (γ) of tests that must produce a 3- to 5-log_10_ reduction. A value of 95% is reasonable based on statistical norms.

In Parker’s [21] review, the reproducibility standard deviations for the HSCT and the UDM with *P. aeruginosa* did not exceed 0.8 and 1.1 log_10_, respectively. The HSCT data was from multi-laboratory collaborative studies that included four different disinfectant products [24, 25]. The UDM data was from a multi-laboratory studies of single quaternary ammonium disinfectant product [26].

Because our study only included a quaternary ammonium chloride product, our findings should not be generalized to apply to disinfectant groups with dissimilar modes of action such as chlorinated compounds, per-oxygen compounds, and organic acids. Accordingly, our findings only apply to *P. aeruginosa.* Despite these limitations, our results are important because researchers, product developers, and regulators need standard methods to be reproducible across a broad range of disinfectants and microbes (including quaternary ammonium chloride products and *P. aeruginosa*).

The results in this paper support a need for deeper analysis of testing materials and procedures as a potential cause of the problem. Thus, we recommend the QM be subjected to ruggedness testing to assess its sensitivity to small changes of operational factors such as inoculum preparation, soil preparation, inoculum drying, carrier surface preparation, and recovery procedures. It is our expectation that modest changes to the QM will remove much of the variability we observed. Ruggedness testing is a critical aspect of disinfectant efficacy test method development though it is often overlooked.

### Final thoughts and recommendation

The reproducibility studies shared in this paper were conducted after two years of intensive use of, and improvements to, the QM with a goal of assuring reasonable alignment between labs. The analysts in the EWG labs have had hundreds of hours of hands-on experience prior to this study and have cross-trained each other on all aspects of the QM. The alignment of outcomes achieved when analysts from different laboratories performed the same test in the same laboratory confirm that reproducibility problems were not due to analyst proficiency issues. A strong research partnership across government, academic and industry experts is recommended to solve the reproducibility problem with the QM. Solving the reproducibility problem would then enable risk-benefit analyses related to test method correlation.

## Acknowledgements

Our appreciation to the members of the Efficacy Working Group (EWG) and Dr. Florian Brill at The Institute for Hygiene and Microbiology. The EWG is a consortium consisting of Arxada LLC, The Clorox Company, Ecolab Inc., Procter & Gamble, Pilot Chemical, Reckitt, and Stepan Company. The EWG is administered by Patrick Quinn of The Accord Group and Rhonda Jones RM(NRM) of Scientific & Regulatory Consultants, Inc.

## Notes

### Competing Interest Statement

The authors have declared no competing interest.

